# Mosquito diversity and arbovirus circulation at the Taï National Park, western Côte d’Ivoire

**DOI:** 10.1101/2025.07.10.664066

**Authors:** Silvan Hälg, Zonzéréké Coulibaly, Christian Beuret, Julien Z. B. Zahouli, Pie Müller

## Abstract

Understanding the ecological and virological dynamics of mosquito populations across different landscapes is essential for predicting and mitigating the risk of arbovirus spillover. From December 2021 to September 2022, we conducted an entomological and molecular survey across sylvatic, transition, and urban zones in the Taï region of western Côte d’Ivoire. We collected 4,414 mosquitoes, representing 29 taxa, and grouped them into 1,553 pools of the same taxa for virus detection and identification. While *Eretmapoditis* sp. were dominant across all zones, their species composition varied between the zones, with *Er. fraseri* being most abundant in the sylvatic and urban zones. In the transition zone *Er. quinquevittatus* was the dominant species. Abundance of other species also varied across zones. *Aedes aegypti* and *Anopheles* sp. were most frequent in the urban zone with *An. gambiae* s.l. and *An. paludis* being the most common ones. Molecular screening revealed flavivirus RNA being present in multiple pools, with the highest detection rates in the transition zone. We were able to assemble three flavivirus contigs, including Cimo flaviviruses II and VIII, known as insect-specific flaviviruses that can influence vector competence. Notably, *Anopheles* sp. had the highest estimated infection rates in the sylvatic and transition zones, suggesting a potential role in arbovirus ecology. Our findings suggest that *Eretmapodites* sp., particularly *Er. fraseri*, may serve as key bridge vectors due to their widespread presence, mammalian feeding behaviour and potential virus competence. The transition zone emerged as a hotspot for arbovirus diversity and vector-host interactions. This study provides the first comprehensive ecological and molecular characterization of mosquito communities in the Taï region since more than twenty years and underscores the importance of longitudinal and habitat-specific surveillance for anticipating spillover risks.

**Author summary:** Mosquitoes are important to study because they can carry viruses that spread to humans and animals. These viruses, known as arboviruses, often come from wildlife and can spill over into people, especially when natural and human environments overlap. From late 2021 to 2022, we collected and studied mosquitoes in western Côte d’Ivoire, across forest, village-forest edges, and towns. We wanted to understand which mosquito species were present, how their populations varied across different landscapes, and whether they carried any viruses. We found that one group of mosquitoes, called *Eretmapodites*, was common in all areas, with different species dominating in different zones. We also found that some mosquito groups, including *Anopheles*, carried viruses that could potentially affect humans or influence how diseases spread. The area between forest and town, the transition zone, showed the greatest variety of viruses and mosquitoes. Our work helps show how changes in land use and mosquito ecology can affect the risk of disease outbreaks. By understanding where and when mosquitoes are most likely to carry viruses, we can improve surveillance and better prepare for future disease emergence. This is especially important in regions where people and wildlife live close together.

## Introduction

Arboviruses such as dengue virus (DENV), yellow fever virus (YFV), chikungunya virus (CHIKV), Zika virus (ZIKV) and West Nile virus (WNV) pose a significant public health threat in Africa, putting more than 831 million people at risk (about 70% of the continent’s population) (1, 2). The threat of arboviral outbreaks is exacerbated by rapid urbanisation, increased human mobility and ecological disturbance, all of which facilitate the spread of viruses and their mosquito vectors (3). Despite the existence of surveillance, prevention and control programmes, countries like Côte d’Ivoire continue to experience recurrent arboviral disease outbreaks, both in urban and rural settings (4, 5).

Over the past decade, Côte d’Ivoire has faced several outbreaks of dengue and yellow fever. Notably, in 2017, the country reported 623 suspected dengue cases (192 confirmed, two fatalities) and 68 suspected yellow fever cases (one confirmed) (4). A more severe dengue outbreak followed in 2019, with 3,201 suspected cases, 281 confirmed, and two deaths, in addition to 89 confirmed yellow fever cases and one fatality (5). Most recently, dengue outbreaks were reported between April and September 2024, with cases recorded across hospitals nationwide (5). However, due to frequent underreporting and the frequent misdiagnosis of arboviral infections as malaria, actual case numbers are likely significantly higher (6, 7).

Arboviruses are typically maintained in two ecologically distinct transmission cycles: the sylvatic-enzootic cycle, involving wildlife and forest-dwelling mosquitoes and the epidemic-urban cycle, which involves human-to-human transmission via anthropophilic mosquitoes such as *Aedes aegypti* and *Aedes albopictus* (8–12). Mosquitoes feeding on both wildlife and humans, so-called ‘bridge vectors’, may facilitate viral spillover events (13, 14). The likelihood of such events depends not only on mosquito abundance and species distribution, which are influenced by climate, habitat and breeding site availability, but also on factors such as mosquito survival, feeding preferences, host availability and vector competence (15).

Côte d’Ivoire harbours a rich and diverse mosquito fauna, including genera such as *Aedes*, *Anopheles*, *Culex*, *Eretmapodites* and *Mansonia*, present across sylvatic, rural and urban environments (16, 17). While *Aedes* and *Culex* are recognized as major arbovirus vectors, other genera such as *Eretmapodites*, typically associated with forested habitats and tree holes, have also been found to harbour arboviruses, suggesting a potential role as secondary or bridge vectors (8, 13, 18). Their ecological flexibility and feeding behaviour, including occasional anthropophagy, may enable these mosquitoes to mediate cross-species virus transmission [17].

A notable ecological and epidemiological hotspot in Côte d’Ivoire is the Taï National Park, located in the southwest of the country. As one of the largest remaining tracts of primary tropical rainforest in West Africa, and as a UNESCO World Heritage Site, the park is renowned for its exceptional biodiversity and ecological significance (19, 20). The park also harbours a highly diverse mosquito community, including 12 genera: *Aedes*, *Anopheles*, *Culex*, *Uranotaenia*, *Aedomyia*, *Coquilletidia*, *Culiseta*, *Eretmapodites*, *Harpagomyia*, *Mansonia* and *Toxorhynchitinae* (21–23).

Previous studies in the Taï region have identified several viruses in local mosquito populations, such as Nounané virus, Moussa virus, Cavally virus, Gouléako virus, Herbert virus and Taï Forest alphavirus, primarily associated with *Culex* and *Uranotaenia* species (21–24). However, no evidence of circulation of major human-pathogenic arboviruses like DENV, YFV or WNV were found, likely due to the timing of mosquito collection during the dry and early rainy seasons (February to June), whereas these viruses tend to circulate later in the rainy season (22, 25–28).

Ecological disturbances, such as deforestation and habitat fragmentation, further complicate the dynamics of arbovirus transmission. Research has shown that mosquito and virus diversity tend to be highest in undisturbed or moderately disturbed habitats (23). Changes in mosquito community composition, such as increased abundance of disturbance-resilient species like *Culex nebulosus*, can lead to higher overall virus prevalence, not necessarily due to increased infection rates, but because of shifts in vector dominance, a phenomenon known as the ‘abundance effect’ (23). Importantly, several of the viruses identified in the Taï region belong to the category of insect-specific viruses (ISVs), viruses that naturally infect mosquitoes but are incapable of replicating in vertebrate hosts (29). These ISVs are of growing interest in arbovirology, as they may play a critical role in modulating arbovirus transmission (30–32). Through mechanisms such as superinfection exclusion, ISVs can interfere with the replication and dissemination of medically important arboviruses in mosquito vectors, potentially reducing their transmission efficiency (33). Moreover, the evolutionary relationships between ISVs and vertebrate-pathogenic arboviruses suggest that ISVs may serve as evolutionary precursors and reservoirs of genetic diversity (34–36). Their biological safety profile and natural restriction to invertebrate hosts also make ISVs attractive candidates for developing novel vaccine platforms (37, 38) and biocontrol tools (39, 40). As such, understanding ISV, arbovirus interactions within natural mosquito populations holds considerable promise for future outbreak mitigation and vaccine research in arbovirus-endemic regions such as Côte d’Ivoire

Taken together, these findings underscore the importance of understanding how mosquito diversity, habitat disturbance and ecological interfaces influence arbovirus ecology and emergence. As anthropogenic activities continue to alter forest landscapes, the need for integrated entomological and virological surveillance becomes more urgent.

In this study, we aimed to extend research further back in time by conducting a comprehensive mosquito survey in and around the Taï National Park across nearly a full seasonal cycle, covering undisturbed forest, transitional and urban landscapes. Our objectives were to characterise the mosquito community composition, identify potential bridge vector species and assess their role in arbovirus and ISV transmission dynamics in one of West Africa’s most biodiverse and epidemiologically significant ecosystems.

## Methods

### Study area

We collected mosquitoes from three distinct locations representing the ‘sylvatic’, ‘transition’ and ‘urban’ zones in and around the Taï National Park in western Côte d’Ivoire (Figure 1). Taï National Park is one of the few remaining primary rainforest areas in West Africa, spanning nearly 600,000 ha. It hosts over 1,800 plant species, 54% of which are exclusive to the West African forest flora, with 138 endemic species only occurring in the Taï National Park (41). The forest also hosts a remarkable diversity of wildlife, making it one of the most important biodiversity hotspots in West Africa. The park is home for over 250 bird species and 47 mammal species including 11 species of primates such as chimpanzees (42). The area surrounding the park (Figure 1) is dominated by agricultural activities, including land development, lowland cultivation and pond creation for fish farming and human settlements, which have contributed to increased mosquito densities (43).

**Figure 1:**
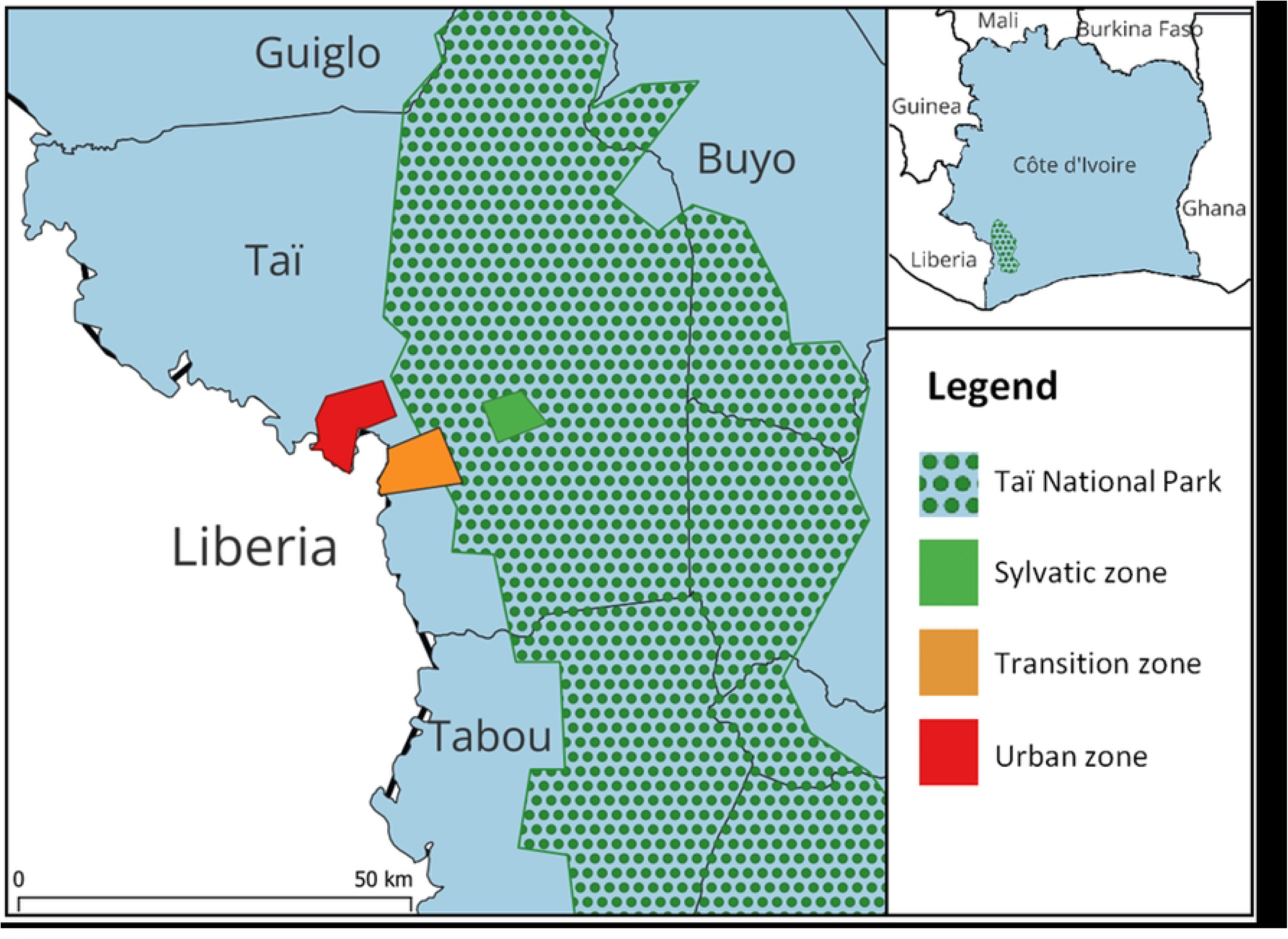
Map of the study area located in and around the Taï National Park, western Côte d’Ivoire. Mosquitoes were collected in the sylvatic, transition and urban zones. The map was created using QGIS 3.34.13 (44).

For the sylvatic zone, we chose trap sites inside the primary forest around the Station de Recherche Ecologique de Taï (SRET; 5.833373 N, 7.342714 W). In the transition zone, we placed the traps in the village of Pauléoula (5.824167 N, 7.401908 W) and its adjacent rubber plantations located at the forest fringe. For the urban zone, we selected the town of Taï (5.874790 N, −7.456201W) that is slightly more distant than Pauléoula from the national park. Taï shows a more urbanised character with improved houses and is more densely populated than Pauléoula.

The climate in the study area is sub-equatorial, hot and humid throughout the year, with an average annual rainfall exceeding 1,600 mm across the entire massif and an average temperature of about 26 °C (45). A year has typically four distinct seasons: a long rainy season from March or April to July, a short dry season in August, a short rainy season in September and October, and a long dry season from November to February or March.

### Meteorological data

To collect actual meteorological data during the study, including relative humidity, temperature and precipitation, we installed a Vantage Pro 2 weather station (Davis instruments, Hayward, CA, USA) at each mosquito-sampling site, relocated the station accordingly as sampling locations changed.

### Mosquito sampling

From December 2021 to September 2022, we collected adult mosquitoes using Centres of Disease and Prevention (CDC) miniature light traps (model 512) and box gravid traps sourced from BioQuip (BioQuip Products, Inc., Rancho Dominguet, CA, USA). At a time, we placed four traps of each type in one of the three zones (sylvatic, transition and urban) for five consecutive days (240 h). We then rotated the traps to the next zone, continuing until we had sampled each zone eight times. Within a zone, we determined the trap position randomly using QGIS Version 3.22.0 (44).

Due to the difference in the landscape of the three zones, we adjusted the placement of light traps accordingly. In the sylvatic zone, we deployed two light traps in the canopy, at a height of 10-15 m above ground, and two traps at 1.5 m above ground. To place the traps in the canopy, we used a slingshot (Bugnard AG, Zürich, Switzerland) to shoot a weight attached to a rope over a branch. Then we hoisted the traps into position with a belay attached to the end of the rope. In the transition zone, we placed two light traps in the rubber plantation adjacent to the village Pauléoula and two traps near houses within the village. Unlike in the sylvatic zone, in the transition zone, as well as in the urban zone, we attached the four light traps to trees at 1.5 m above ground.

As an attractant, we used CO_2_ produced by sugar-yeast solutions contained in bottles and released via tubes connected to the trap cover (Figure 2a). We prepared three solutions with varying concentrations: 500 g sugar and 21 g yeast in 1.5 l water (high concentration), 250 g sugar and 14 g yeast in 1.5 l water (medium concentration), and 250 g sugar and 7 g yeast in 1.5 l water (low concentration). Each trap was fitted with three bottles containing one of each concentrations, ensuring a steady production of CO_2_ over 12 h as described in Steiger et al. (46). We replaced the yeast solutions every 12 h and changed daily the containers with the caught mosquitoes.

**Figure 2:**
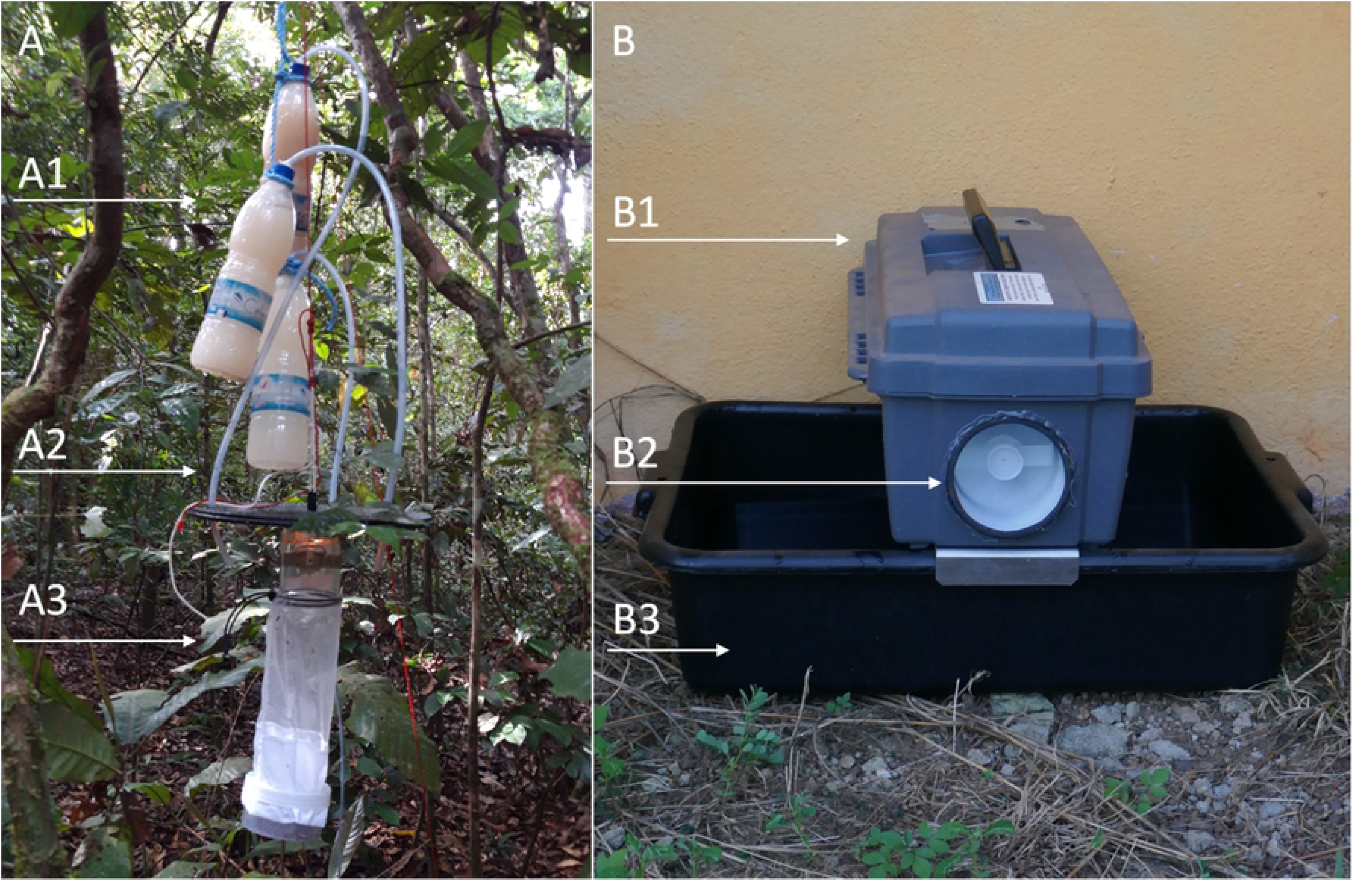
Trap types used to collect mosquitoes in three ecological zones in the Taï area, western Côte d’Ivoire. A CDC miniature light trap (A3). Mosquitoes were attracted by the CO2 produced by aqueous sugar-yeast solutions contained in three bottles that were positioned above the trap (A1) and connected via tubes (A2). Each solution had a different sugar-yeast concentration (low, medium and high), resulting in staggered CO_2_ peak production times, thereby ensuring a steady flow of CO_2_ over time. B Box gravid trap. The trap attracts gravid female mosquitoes, seeking an ovipositon site, by water treated with a standardised hey infusion (B3). An electric fan (B2) then sucked the mosquitoes into a container (B1).

**Figure 3:**
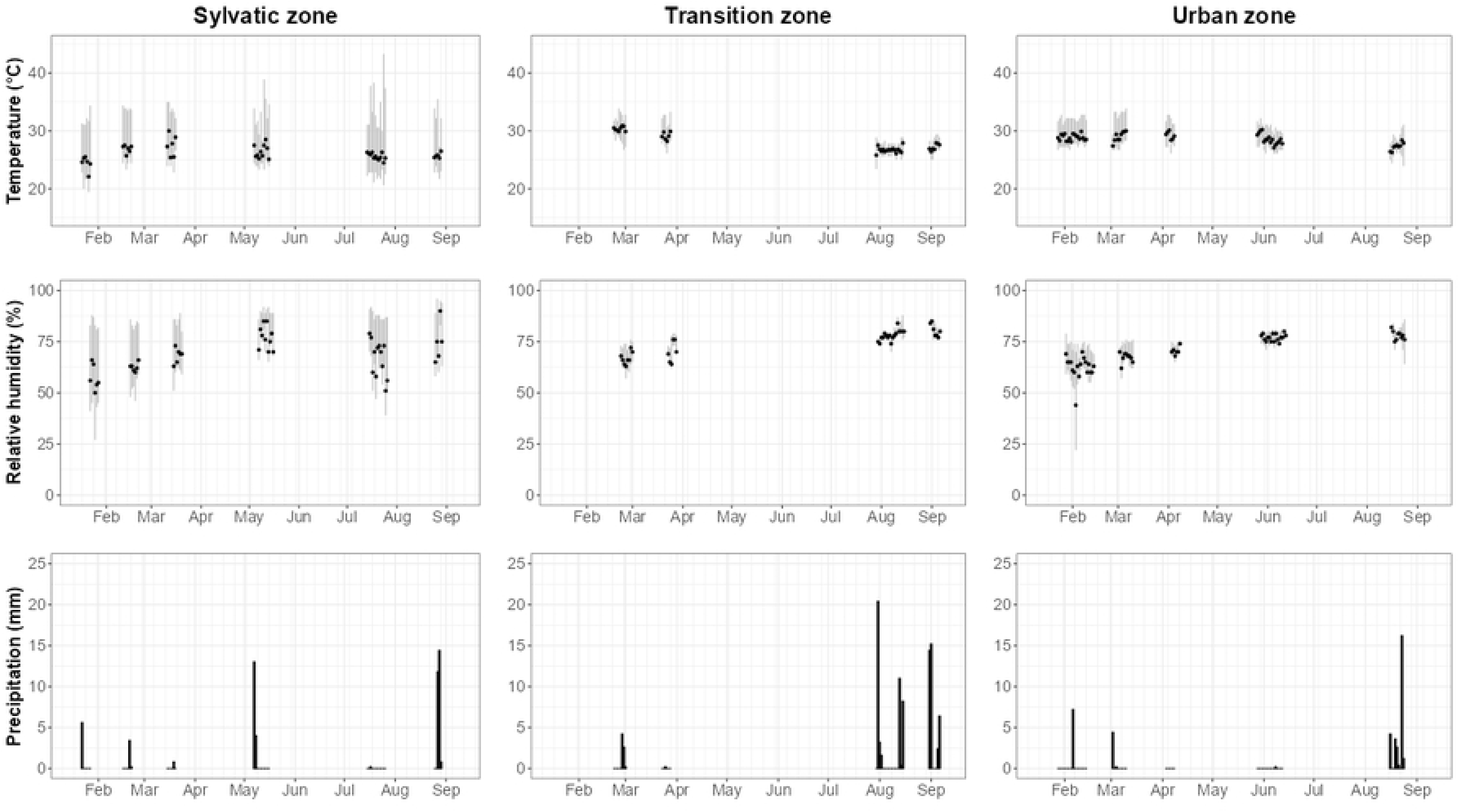
Meteorological data recorded during the mosquito collections using a mobile weather station in

We placed the box gravid traps strategically to protect them from rain. In the sylvatic zone, we positioned the traps under trees and roots. In the urban zone, we placed the traps under roof overhangs or small shacks. In the transition zone, we placed two box gravid traps under roof overhangs and two on the ground in the rubber plantations beneath the shelter of a tree or bush. On the first day of setting up at a new location, we filled each trap’s container with 5 l water and added one Synthetic Hay-Infusion *Culex* Lure (BioQuip). We then left the container unchanged for the following five days, while every 24 h we replaced the collection box containing the trapped mosquitoes. In addition, we collected any visible mosquito egg rafts and juvenile stages from the water containers and transported them together with the adults to a temporarily set up field laboratory in Taï.

### Mosquito identification

We transferred the larvae and eggs from the box gravid traps into plastic cups and fed them with yeast every two days until pupation, changing the water every second day. We anesthetised adult mosquitoes, whether collected from the field or reared in the field insectary, with 100% chloroform for subsequent morphological identification. For identification, we sorted specimens in Petri dishes and classified them by sex, genus and the lowest possible taxonomic level with a stereo microscope. We used the morphological identification keys of Mattingly (47) for culicine and the key by Gilles and De Meillon for anopheline species (48). Mosquitoes were pooled into groups of up to ten individuals, matched by species and sex.

### Mosquito preservation and RNA extraction

After identification, mosquitoes were briefly immersed in 100% ethanol to increase tissue permeability, then transferred into an RNA-preserving solution, prepared according to (49), and stored them overnight at 4 °C. The following day, samples were removed from the solution, air-dried, transferred into fresh 1.5 ml Eppendorf tubes in pools of 10, separated by taxa and site, and stored at –20 °C on site before they kept at −80 °C for longer-term storage.

For homogenising the mosquito samples, we transferred the mosquito pools to 96-well collection plates. To each well, we added 600 µl of Inhibitex Buffer (Qiagen, Hilden, Germany) and a 3 mm tungsten carbide bead (Qiagen). Using a Qiagen TissueLyser Universal Laboratory Mixer-Mill Disruptor, we homogenised the mosquitoes for 3 min at 30 Hz, rotated the plates by 180° and repeated the process to ensure uniform homogenisation. Finally, we centrifuged the plates at 3,220 × g for 5 min.

For virus inactivation, we combined 100 µl of each homogenate with 400 µl of AVL buffer directly in a MagNA Pure 96 Processing Cartridge (Roche, Basel, Switzerland) and stored the remaining homogenates at –80 °C for future analyses. Each processing plate included two positive control wells: one containing inactivated yellow fever virus 17D, and the other containing chikungunya virus. Total nucleic acid was extracted using the MagNA Pure DNA Viral NA LV 2.0 Kit (Roche, Switzerland) on the MagNA Pure 96 System (CE-IVD, Roche)

### Detection of viral RNA

To screen the mosquito pools for viral RNA, we used reverse transcription PCR (RT-PCR) with separate PCR protocols for alphaviruses and flaviviruses.

For the detection of alphaviruses, we ran PCR reactions in volumes of 25 µl using the forward and revers primers, VIR966F and VIR966R (Table 1) (50). The reactions included 5 µl RNase-free H₂O, 1 µl dNTPs (10 mM), 1.5 µl of each primer (10 µM), 5 µl Q-Solution, 5 µl 5× OneStep RT-PCR Buffer, 1 µl OneStep RT-PCR Enzyme Mix (Qiagen) and 5 µl template RNA. Cycling conditions were as follows: reverse transcription at 50 °C for 30 min; initial denaturation at 95 °C for 15 min; 45 cycles of denaturation at 94 °C for 20 s annealing at 55 °C for 20 s and extension at 72 °C for 20 s.

**Table 1:**
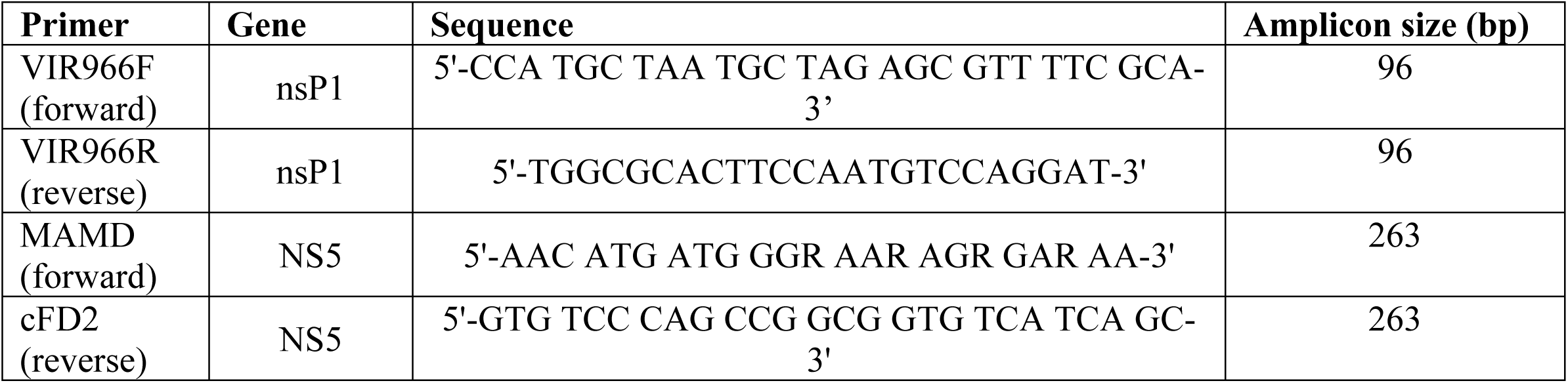
Primers used in the RT-PCR reactions for the identification of alpha- and flaviviruses.

For the detection of flaviviruses, we ran PCR reactions in volumes of 50 µl using the forward and reverse primers, MAMD and cFD2 (Table 1) (51). The reaction mix consisted of 25 µl RNase-free H₂O, 2 µl dNTPs (10 mM), 3 µl of each primer (10 µM), 10 µl 5× OneStep RT-PCR Buffer, 2 µl OneStep RT-PCR Enzyme Mix (Qiagen),and 5 µl template RNA. Cycling conditions were as follows: reverse transcription at 50 °C for 30 min; initial denaturation at 95 °C for 15 min; 45 cycles of denaturation at 94 °C for 1 min, annealing at 50 °C for 1 min; extension at 72 °C for 1 min; amd a final extension at 72 °C for 7 min.

To visualise the amplicons from both assays, we ran the PCR products using the QIAxcel DNA Screening Kit in combination with a QIAxcel System (Qiagen). For visualisation and data interpretation, we employed the QIAxcel ScreenGel Software (Qiagen). We then selected those amplicons that were tested positive for either an alpha- or a flavivirus for Sanger amplicon sequencing.

In preparation for the sequencing, we purified the PCR amplicons using the QIAquick PCR Purification Kit (Qiagen), following the manufacturer’s protocol. For each primer, VIR966F, VIR966R, MAMD and cFD2, we carried out a separate sequencing reaction using the corresponding PCR product as a template. Each 20 µl sequencing reaction contained 4 µl BigDye Terminator v3.1 Ready Reaction Mix, 2 µl 5× BigDye Sequencing Buffer (Thermo Fisher Scientific, Waltham, MA, USA), 2 µl primer (3.2 µM), 2 µl purified PCR fragment and 10 µl RNase-free water. Thermal cycling conditions were as follows: initial denaturation at 95 °C for 1 min; followed by 25 cycles of denaturation at 95 °C for 10 s, annealing at 50 °C for 5 s and extension at 60 °C for 2 min.

We cleaned the sequencing reactions using DyeEx 2.0 Spin Columns (Qiagen), according to the user manual, and performed capillary sequencing on ABI Prism® 3130 and 3130xl Genetic Analyzers using the SeqStudio Genetic Analyzer Cartridge v2 (Thermo Fisher Scientific).

### Data analysis

We performed all statistical analysis in R version 4.1.3 (52). A species rarefaction curves was done using the vegan package (53). We calculated summary statistics, including the relative abundance (F) and frequency of occurrence (C) of each mosquito taxon (species or genus) in each zone across both trap types. To test whether taxon abundance across zones differs, we performed a *χ^2^*-test with Monte Carlo simulation (54), combining the numbers of individuals from the rare taxa with an overall relative abundance (F) of less than 5% (see S1 Additional file, Table A1). We then investigated species similarity between the three zones by calculating the Sorenson’s coefficient (*S*) (55).

We calculated the absolute abundance of each taxon by summing up all the collected individuals for each taxon. The formula for the relative abundance of each taxon (*F*), a measure of how common a species is compared to others in a defined location (56), is as follows (57): 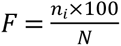, where *n*_i_ is the number of individuals of species *i*, and *N* is the total number of individuals of all species.

For each mosquito species and sampling location, we also calculated the frequency of occurrence (*C*) as: 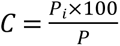, where *P*_i_ is the number of surveys containing the species, and *P* is the total number of surveys conducted.

The Sorenson’s coefficient was calculated as follows: 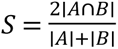, where *A* and *B* are the mosquito species found in the different zones, ∣*A*∩*B*∣ is the number of shared mosquito species between the two compared zones and ∣*A*∣ and ∣*B*∣ are the number of species found in each zone. The coefficient ranges from 0 (no similarity) to 1 (perfect similarity).

Sequence data were aligned into contigs using CodonCode Aligner software, version 8.0.2. Each forward and reverse sequence as well as contig was identified using online BLAST (58). The infection rate was calculated using a maximum likelihood estimation (MLE) to account for the variable pool sizes. It was calculated in Python version 3.12 using the libraries ‘pandas’ (59), ‘numpy’ (60) and ‘scipy.optimize’ (61). The formula to calculate the MLE is as follows: 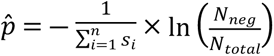, where *p̂* is the estimated infection rate (MLE), *s_i_* is the size of the *i*-th pool, *N_neg_* is the number of negative pools and *N_total_* is the total pool number. We calculated the MLE at the genus level, and not at the species level, because for some species we only had a few pools to work with. Also, were not able to identify all positive pools to species level.

## Results

### Meteorological data

Meteorological data, including daily maximum, minimum, and mean temperatures, relative humidity, and rainfall, were recorded and are summarized in table 2. While all three zones, sylvatic, transition, and urban, experienced broadly similar seasonal patterns, distinct differences were observed in their thermal and hydrometeorological profiles.

**Table 2.** Summary of meteorological conditions across sylvatic, transition, and urban zones.

|  | **Mean temperature**<br>(temperature range) | **Relative humidity**<br>(humidity range) | **Total rainfall, rainy days**<br>(maximum daily rainfall) |
| --- | --- | --- | --- |
| **Sylvatic zone** | **23.2 °C**<br>(19.4 °C – 26.6 °C) | **68.3%**<br>(50% - 90%) | **54.2 mm, 45 days**<br>(14.4 mm, 25. Aug. 2022) |
| **Transition zone** | **26.5 °C**<br>(24.5 °C – 29.4 °C) | **74%**<br>63% - 85%) | **90.4 mm, 38 days**<br>(20.4 mm, 31. July 2022) |
| **Urban zone** | **27.0 °C** | **70.4%** | **40.2 mm, 58 days** |
|  | (23.9 °C – 28.9 °C) | (44% - 82%) | (16.2 mm, 23. Aug. 2022) |

The sylvatic zone exhibited the lowest overall mean temperature (23.2 °C), with daily means ranging from 19.4 °C to 26.6 °C. In contrast, the transition and urban zones were warmer, averaging 26.5 °C and 27.0 °C, respectively. Temperature ranges were slightly broader in the transition zone than in the urban zone.

Relative humidity was highest in the transition zone (mean 74%, range 63–85%), followed by the urban zone (70.4%, range 44–82%), and lowest in the sylvatic zone (68.3%, range 50–90%). Despite its lower average humidity, the sylvatic zone showed the widest range in values.

Rainfall patterns also varied: the transition zone recorded the highest total rainfall (90.4 mm over 38 days), followed by the sylvatic zone (54.2 mm over 45 days), while the urban zone experienced the lowest rainfall (40.2 mm over 58 days). The maximum daily rainfall was highest in the transition zone (20.4 mm), compared to 16.2 mm in the urban zone and 14.4 mm in the sylvatic zone.

### Mosquito species composition and frequency

In total, we collected 4,603 mosquito specimens with 2,192 (1,515 females), 1,185 (1,091 females) and 1,226 (983 females) specimens from the sylvatic, transition and urban zones, respectively. We identified all specimens at least to genus level and found eight genera across the three zones with varying proportions across the zones: *Aedes*, *Anopheles*, *Coquillettidia*, *Culex*, *Eretmapodites*, *Mansonia*, *Toxorhynchites* and *Uranotaenia* (Figure 4 and S1 Additional file, Table A2). The most common genus across all zones was *Eretmapodites*. In contrast, the second most common genus differed across zones: *Uranotaenia* was predominant in the sylvatic zone, *Mansonia* in the transition zone and *Aedes* in the urban zone.

**Figure 4:**
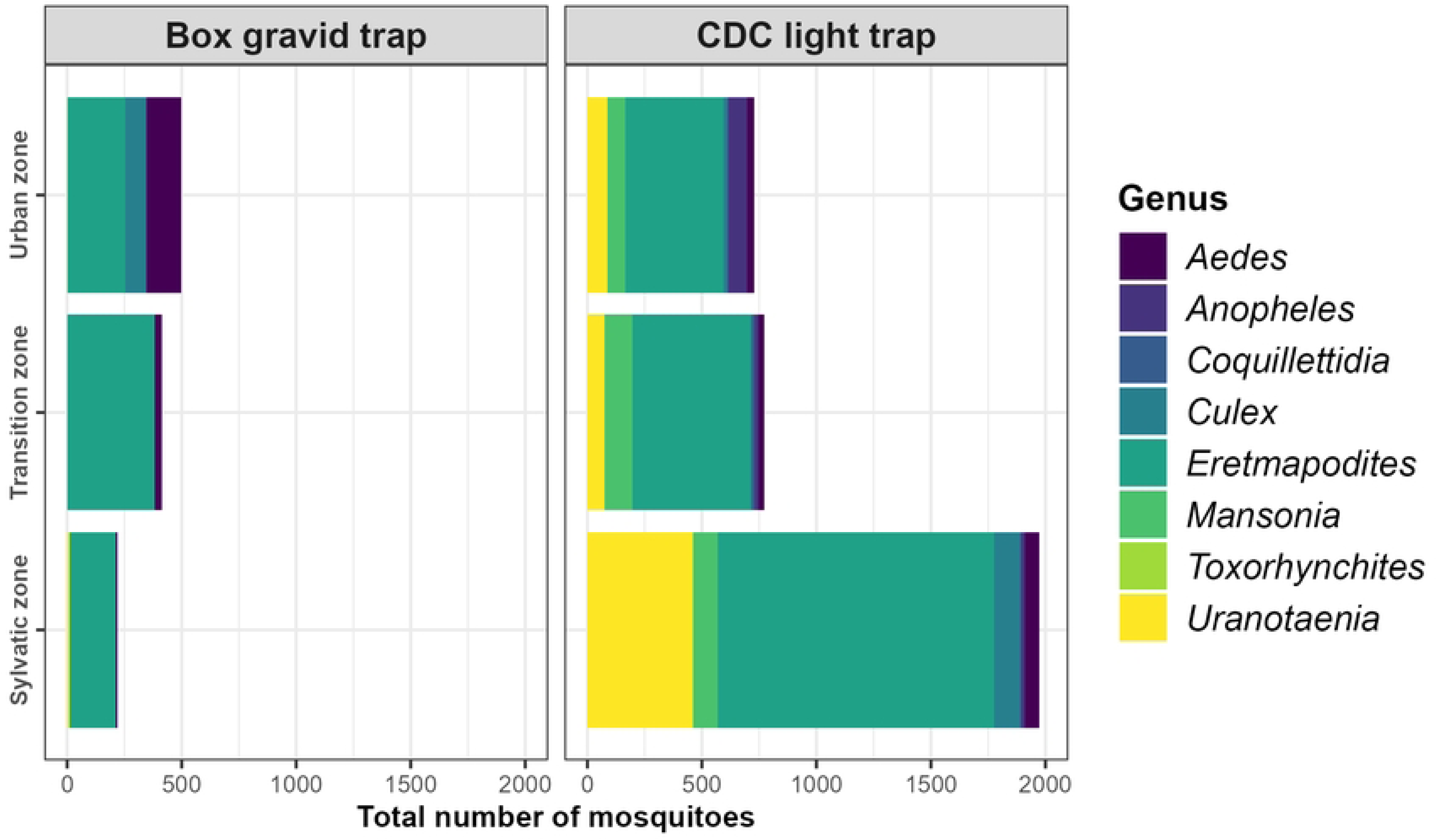
Composition and frequency of mosquito genera across the sylvatic, transition and urban zones by trap type in the Taï area, western Côte d’Ivoire.

Across the three zones, we were able to identify 64.1% of the specimens to species level, resulting in 29 species (S1 Additional file, Table A3). The remaining 35.9% were only identified to genus level. In the sylvatic zone, the most abundant species was *Eretmapodites fraseri* (57.5%), followed by *Uranotaenia bilineata* (25.5%). In the transition zone, *Eretmapodites quinquevittatus* (48.4%) was the most abundant species, followed by *Er. fraseri* (24.6%). *Eretmapodites fraseri* (37.4%) was also the most abundant species in the urban zone, followed by *Ae. aegypti* (27.4%). The species rarefaction curves in Figure 5 show that we collected nearly all different mosquito taxa since they were reaching a plateau as a function of collection effort.

**Figure 5:**
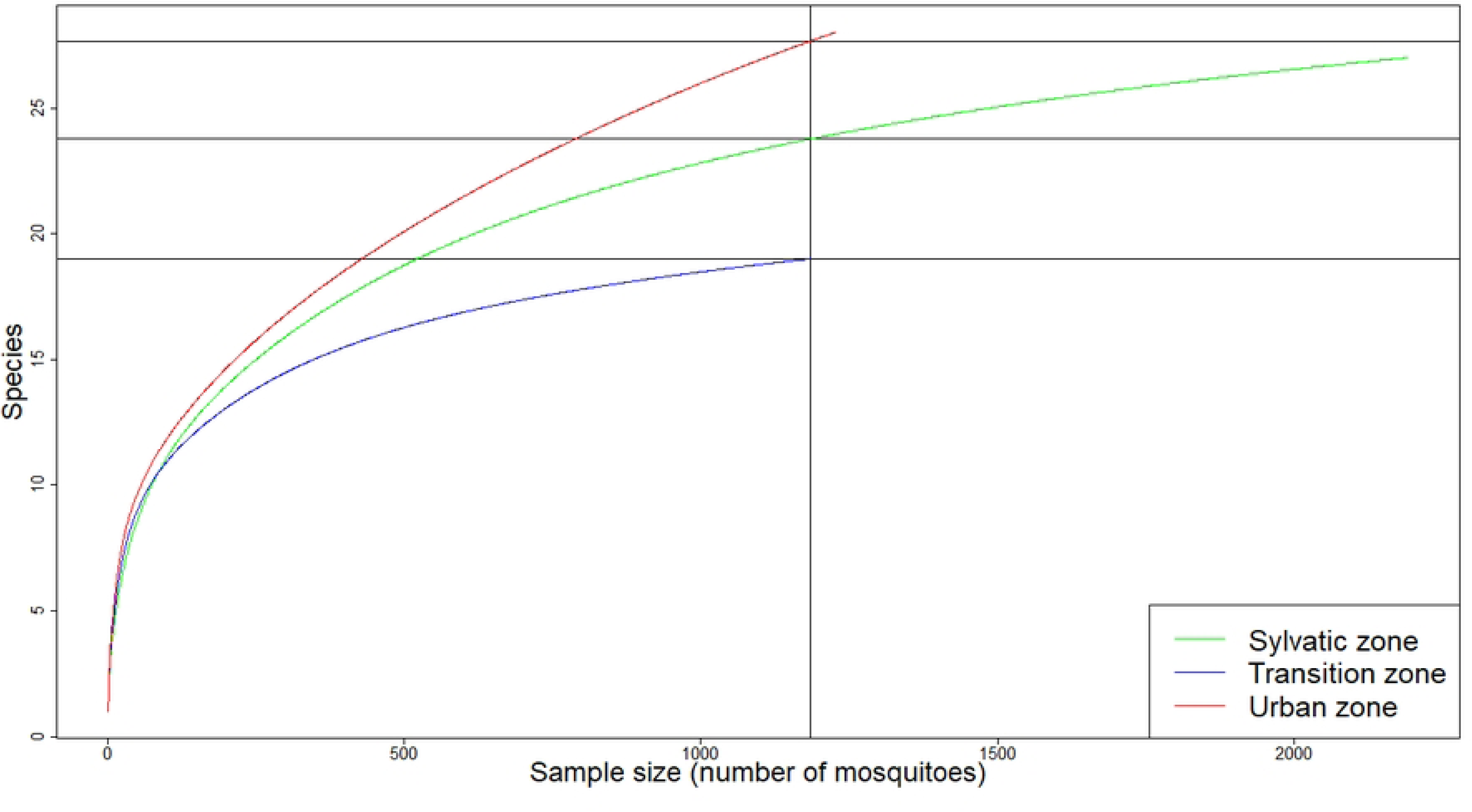
Species rarefaction curves for mosquito taxa in the Taï area, western Côte d’Ivoire. The curves show that most mosquito taxa have been identified across the three zones. The three horizontal lines indicating the maximum species identified for each zone and the horizontal line indicate the minimum number of mosquitoes sampled over all three zones.

We determined the gonotrophic stages of the female mosquitoes and found a higher proportion of gravid females in the urban and transition zones, while most mosquitoes in the forest zone were unfed (Figure 6). In the sylvatic zone, we collected 1,515 female mosquitoes (69%), including 449 gravid, three semi-gravid and 1,063 unfed individuals, 53 of which we reared from larvae. In the transition zone, we collected 1,091 female mosquitoes (92%), comprising 796 gravid, two semi-gravid, and 293 unfed individuals, with 28 reared from larvae. In the urban zone, we collected 982 female mosquitoes (80%), including 626 gravid, one semi-gravid and 355 unfed individuals, 78 of which we reared from larvae.

**Figure 6:**
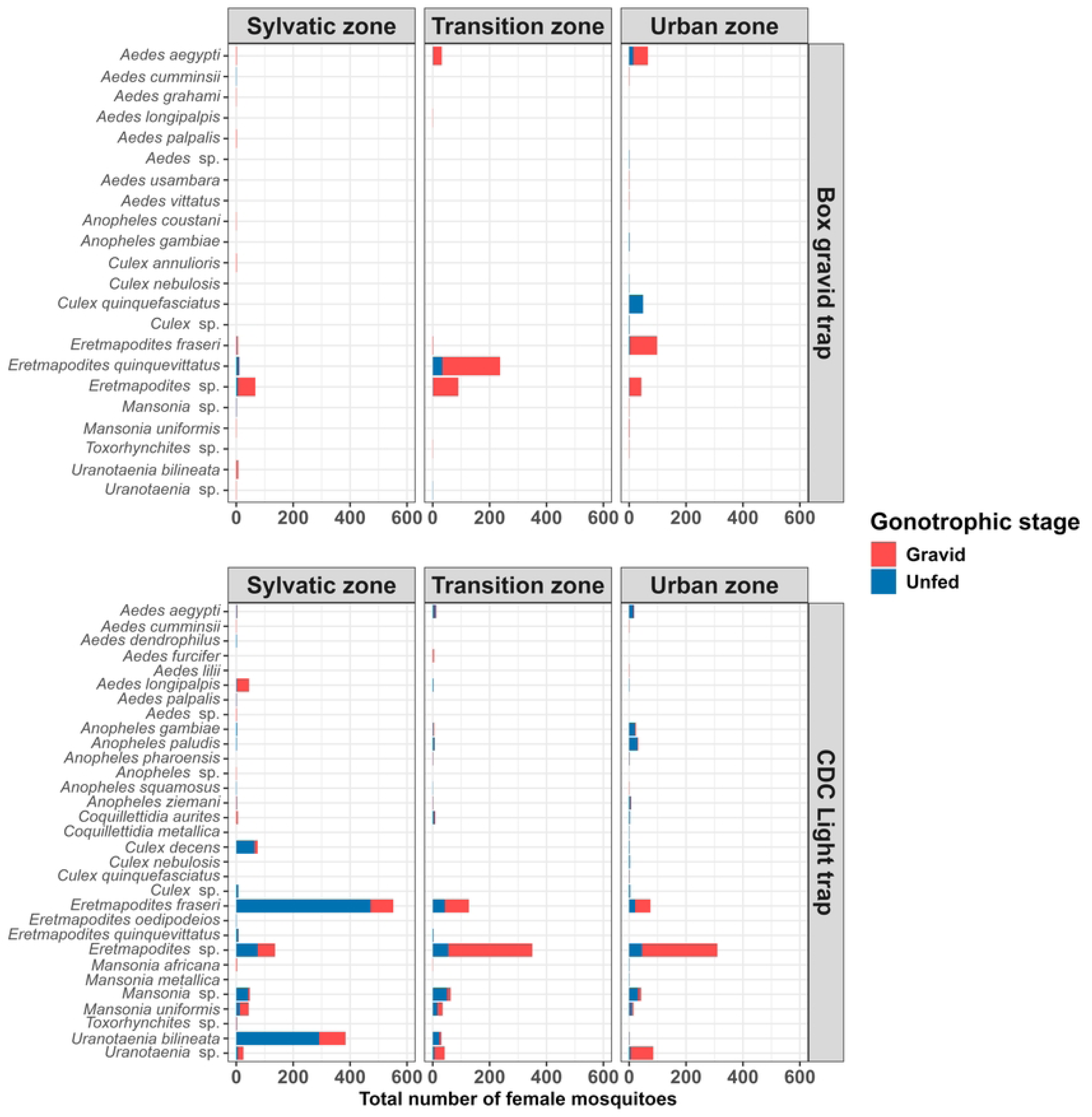
Abundance and gonotrophic stage of female mosquitoes across the sylvatic, transition and urban zones for each trap type in the Taï area, western Côte d’Ivoire. In the urban and transition zones, most of the collected female mosquitoes were gravid, whereas in the sylvatic zone most mosquitoes were unfed.

Each zone shared over 70% similarity in terms of taxa with one of the other two zones (S_sylvatic-transition_ = 0.73; S_sylvatic-urban_ = 0.72; S_transition-urban_ = 0.76). Although we found a high similarity among the zones in terms of shared taxa, their relative abundance differed considerably (*χ^2^* = 225.9, *p* < 0.001; Figure 6). *Aedes* sp., *Culex* sp., *Er. fraseri* and *Ur. bilineata* showed a stronger association with the sylvatic zone, whereas *Coquillettidia* sp. and, especially, *Er. quinquevittatus* were more likely to be found in the transition zone, while *Ae. aegypti*, *Anopheles* sp. and *Cx. quinquefasciatus* were present mostly in the urban zone (Figure 7).

**Figure 7:**
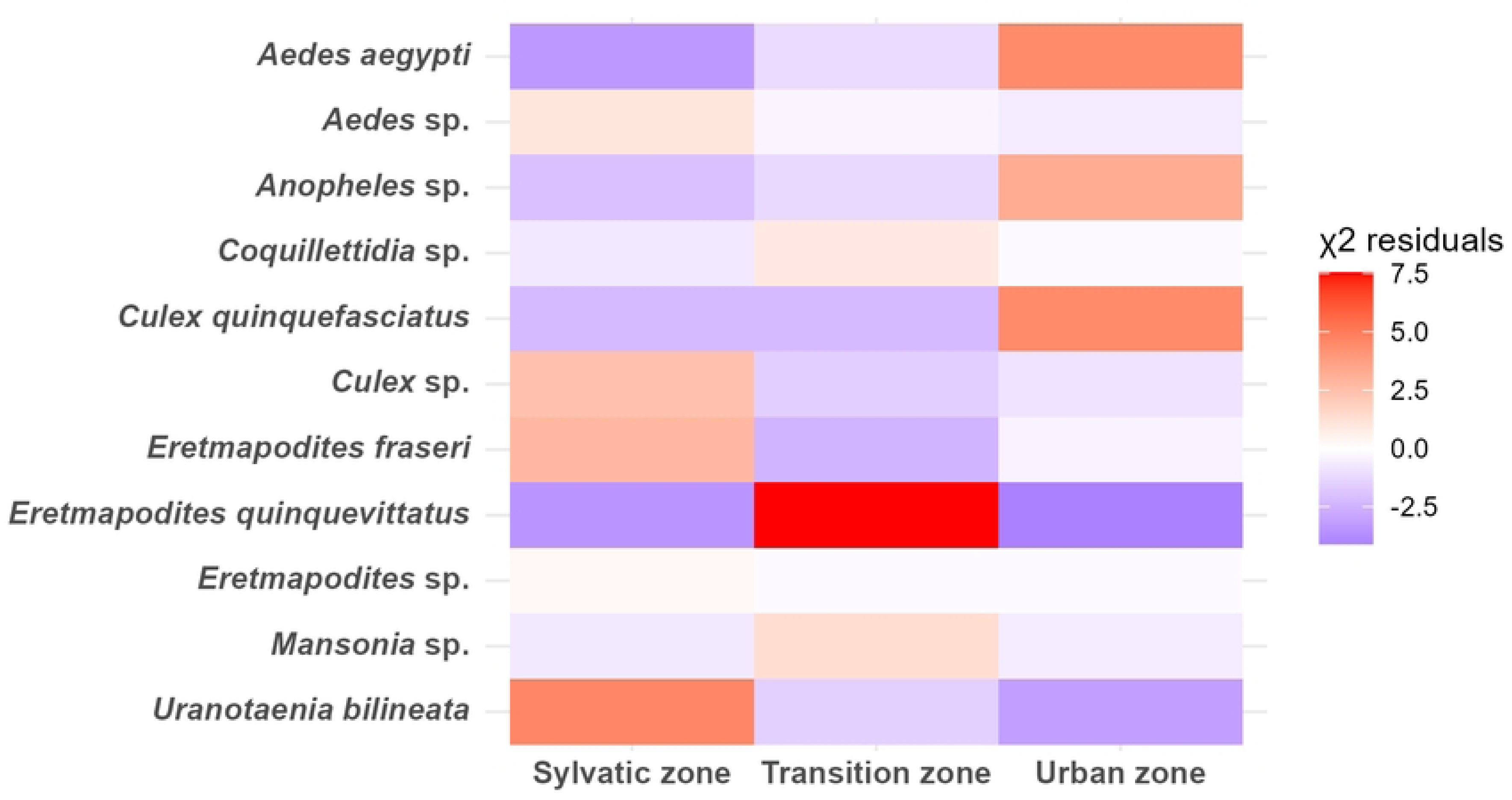
Mosquito abundance across urban, rural and transition zones in Taï, western Côte d’Ivoire. The colour scale represents the magnitude and direction of deviations from a uniform distribution, based on residuals from a *χ²*-test with Monte Carlo simulation. The residuals indicate the difference between the observed and expected frequencies of each mosquito species in each zone with the null hypothesis that the species are evenly distributed and not associated with any particular zone.

### Arbovirus detection

We detected 61 flavivirus-positive pools, 14 in the sylvatic, 35 in the transition and 12 in the urban zones as well as 40 alphavirus-positive pools, 16 in the sylvatic, 13 in the transition and 11 in the urban zones. All flavivirus- and alphavirus-positive samples were sequenced by Sanger sequencing. From the sequencing data, we could assemble three contigs: one from a sylvatic pool and two from transition pools.

Using online BLAST, we analyzed each sequencing read (forward and reverse) as well as each assembled contig individually. In the sylvatic zone, one contig matched an endogenous Culicidae flavivirus (collected on 17 May 2022). In the transition zone, two contigs aligned with Cimo flavivirus II (collected on 28 March 2022) and Cimo flavivirus VIII (collected on 1 May 2022), respectively (see S1 Additional File, Table A5 and S2 Flavivirus.fasta file). No flavivirus-related contigs or BLAST hits were detected in the pools from the urban zone.

BLAST analysis of unassembled reads provided further insights. In the sylvatic zone, we identified one Tembusu virus-like sequence (reverse read, collected on 10 May 2022) and one Anopheles flavivirus-like sequence (forward read, collected on 9 May 2022). In the transition zone, we detected one Usutu virus-like sequence (reverse read, collected on 29 April 2022) and an additional Cimo flavivirus VIII-like sequence (forward read, collected on 2 September 2022). No viral sequences were identified in unassembled reads from the urban zone.

In BLAST analyses of 15 forward primer sequences, we observed alignment with a predicted uncharacterized Aedes aegypti gene (GenBank accession: XR_002500622.1), likely resulting from non-specific primer binding. All BLAST searches for alphavirus-positive pools returned negative results.

In the sylvatic zone, the MLE of infection rates were calculated for three mosquito genera. For *Anopheles*, the infection rate was 5.9% (95% CI: 0.0%–12.1%), based on 17 pools, each comprising a single mosquito. For *Eretmapodites*, the MLE was 0.1% (95% CI: 0.0%–0.4%), derived from 355 pools, with pool sizes ranging from 1 to 12 mosquitoes. For *Mansonia*, the infection rate was 0.9% (95% CI: 0.0%–3.5%), based on 51 pools, with 1 to 10 mosquitoes per pool.

In the transition zone, infection rates were estimated for *Anopheles* and *Eretmapodites*. *Anopheles* showed an infection rate of 6.1% (95% CI: 0.0%–18.1%), calculated from 15 pools, each containing one or two mosquitoes. The infection rate for *Eretmapodites* was 0.3% (95% CI: 0.0%–1.03%), based on 269 pools, with pool sizes ranging from 1 to 11 mosquitoes.

## Discussion

This study provides an integrated ecological and molecular perspective on mosquito diversity and arbovirus circulation across sylvatic, transition and urban landscapes in the Taï region of western Côte d’Ivoire. Over the study period, we identified seven genera within the *Culicinae* subfamily comprising 23 taxa and six *Anopheles* species. While many taxa were the same across landscapes, and species richness was also similar, significant differences emerged in terms of species presence and distribution across the zones.

The key arbovirus vector *Ae. aegypti* was most prevalent in the town of Taï, less common in the transition zone, and nearly absent in the sylvatic zone. This distribution reflects its domestic ecology and presumably its reliance on human-associated breeding sites such as used tyres and water containers (16, 62). Similar patterns were reported in other urbanised regions of Côte d’Ivoire (16, 63). Nevertheless, the mosquito was present across the three landscapes and without knowing its feeding patterns across them we may not exclude its role as a bridge vector.

Intriguingly, *Eretmapodites* were the most frequent mosquito species across all three zones, with *Er. fraseri* dominating the urban and sylvatic zones and *Er. quinquevittatus* the transition zone. The presence of *Eretmapodites* species even in Taï can be attributed to the semi-rural nature of the town, as it is embedded within wetlands and forest fragments and shows features of both traditional and modern human habitation (64). *Eretmapodites* mosquitoes primarily feed on mammals and have been suspected of occasionally biting humans (65), a behaviour that increases their potential as bridge vectors. Indeed, *Er. quinquevittatus* has been shown to be competent for Rift Valley fever virus, while other *Eretmapodites* species have been implicated in the transmission of Spondweni virus and YFV (66–68).

*Anopheles* species were also largely restricted to the urban zone, with *An. gambiae* s.l. and *An. paludis* being the most abundant ones. While *An. gambiae* s.l. is known for its adaptability across both urban and rural settings in Africa, *An. paludis* is not typically associated with anthropogenic environments and is not considered a major vector of human pathogens (69, 70). *Uranotaenia* species, observed primarily in sylvatic and transition zones, are herpetophilic and unlikely to play roles in arbovirus transmission (71).

Consistent with the mosquito species composition, the virological survey also revealed habitat-associated variations in virus detection, with the highest proportion of flavivirus-positive pools occurring in the transition zone. Sequencing confirmed three flavivirus contigs: one from a sylvatic pool matching an endogenous *Culicidae* flavivirus, and two from transition zone pools identified as Cimo flavivirus lineages II and VIII. No confirmed contigs were found in urban pools, despite some PCR-positive results.

The presence of insect-specific flaviviruses (ISFs), particularly in sylvatic and transition zones, mirrors findings from previous studies across Africa and underlines these environments as reservoirs of viral diversity (23, 72–76). Although ISFs are not infectious to vertebrates, they may influence the replication dynamics of pathogenic flaviviruses within mosquitoes and thereby modulate vector competence (73).

The absence of confirmed flavivirus sequences in urban pools may reflect lower viral loads, RNA degradation, or non-specific amplification. This supports the idea that sylvatic and ecotonal zones harbour greater viral diversity due to lower levels of anthropogenic disturbance (23, 77, 78). The detection of numerous virus-positive pools in the transition zone suggests that ecotonal regions, where natural and anthropogenic environments intersect, may facilitate increased virus-vector-host interactions.

Interestingly, *Anopheles* mosquitoes showed the highest MLEs of flavivirus infection in both sylvatic and transition zones. Although *Anopheles* is traditionally associated with malaria transmission, growing evidence indicates its involvement in harbouring other arboviruses, such as WNV and Usutu virus (79–81). These findings reinforce the need to re-evaluate the vectorial capacity of *Anopheles* sp. in arbovirus ecology.

While *Eretmapodites* and *Mansonia* mosquitoes exhibited comparatively lower MLEs, their established or suspected roles in arbovirus transmission, including Rift Valley fever and Semliki Forest virus, highlight their ongoing epidemiological relevance (66–68, 82). However, the wide confidence intervals in MLEs point to the need for larger sample sizes and improved species resolution.

This study has several limitations. First, unusually low rainfall of 184 mm compared to the average of 1,600 mm during the sampling period likely influenced mosquito population dynamics, possibly favouring drought-tolerant species and reducing overall abundance. Additionally, logistical constraints required trap positions remained fixed throughout the study, which may have introduced sampling bias. However, these positions were initially selected randomly, and the consistent performance of CDC trap across the zones suggests that any such bias was minimal. Finally, the use of pooled samples and Sanger sequencing for molecular analysis limited the sensitivity and breadth of virus detection, underscoring the need for high-throughput metagenomic approaches in future studies (83).

Taken together, our findings indicate that *Eretmapodites* species, especially *Er. fraseri*, have strong potential as bridge vectors of arboviruses in the Taï region, given their abundance, mammalian feeding behaviour, and cross-habitat distribution (64–68). The ecological and molecular data converge to suggest that transition zones are critical hotspots for virus emergence and transmission dynamics. The high prevalence of insect-specific flaviviruses, particularly in *Anopheles* mosquitoes, points to an underexplored dimension of arbovirus ecology with implications for vector competence and viral evolution.

This study represents the first comprehensive characterisation of mosquito species diversity and virome composition in the Taï National Park region since over twenty years. Future research should focus on next-generation sequencing, enhanced taxonomic resolution, and longitudinal monitoring of both mosquito populations and their associated viruses. Ecotonal habitats warrant sustained surveillance, as they may act as focal points for spillover events driven by environmental change and human encroachment.

## Acknowledgements

We would like to thank Peho Richard, officer at the Office ivoirien des Parcs et Réserves who connected us with all the persons in Taï. We are greatful to Kara Sepkoh Paulin, head officer of the SRET, and Dr Soro Kafana, researcher at SRET, for facilitating our stay at SRET. We thank our technician Koffi Bernad, who helped us with his expertise in mosquito species identification, and N’guetti Emmanuel for assistance in the field collections.

## Funding

This work was funded and supported by armasuisse W+T, the Swiss Federal Office for Defence Procurement (PN R-3210/045-13).

## Availability of data and materials

Data supporting the conclusions of this article are included within the article and its additional files.

## Author contributions

Conceptualization: SH, CB, JZBZ, PM. Mosquito collection: SH, ZC, JZBZ, PM.

Analysis and interpretation of the data: SH, PM. Writing of the initial manuscript: SH, ZC. Resource coordination: CB, JZBZ, PM. Manuscript revision: SH, ZC, CB, JZBZ, PM. All authors read and approved the final manuscript.

## Ethics approval and consent to participate

Not applicable

## Consent for publication

Not applicable.

## Competing interests

The authors declare that they have no competing interests.

## Supplementary information

### S1 Additional file

**Table A1:** Categories used in the χ2-test with Monte Carlo simulation

**Table A2:** List of all mosquito genera identified in each study zone.

**Table A3:** List of all mosquito species identified in each study zone.

**Table A4:** Total numbers (N), species abundance (F %) and relative abundance (C %) of Culicidae in each study zone.

**Table A5:** Flavivirus positive mosquito pools.

### S2 Flavivirus.fasta

Identified Flavivirus forward, reverse or contig sequences

## Notes

### Competing Interest Statement

The authors have declared no competing interest.

